# Bidirectional modulation of accumbens dopamine by MCH neurons during learning and consummatory behavior

**DOI:** 10.1101/2025.06.05.658170

**Authors:** Liam E. Potter, Brandon A. Toth, Jayeeta Manna, Hannah C. Lyons, Jack R. Evans, Christian R. Burgess

## Abstract

The formation of sensory cue-reward associations is essential for survival, but in the modern calorie-rich and advertising-intensive environment, such associations may become maladaptive - leading to negative health consequences such as obesity or diabetes. Recent research has demonstrated the importance of hypothalamic melanin-concentrating hormone (MCH)-expressing neurons in driving hedonically-motivated feeding and in forming these associations. The MCH system interacts with mesolimbic dopamine (DA) transmission, offering a potential mechanism for the effects of MCH neurons on hedonic feeding and associative conditioning. However, this interaction has not been fully characterized *in vivo* with modern approaches that offer high temporal and spatial resolution. We characterized MCH-DA interactions during feeding and food-motivated Pavlovian conditioning using *in vivo* fiber photometry in the lateral hypothalamus/zona incerta (LH/ZI) and nucleus accumbens (NAc). We found that MCH neuron activity and DA release in the medial-shell of the NAc (mNAcSh) were co-activated during consumption and in response to reward-predicting cues. During consumption, DA release preceded MCH activity, while responses to reward-predicting cues emerged in MCH neurons earlier than in the DA system. Lastly, gain and loss-of function of the MCH system bidirectionally modulates DA release in the mNAcSh. These results indicate that physiological co-activation of the MCH and DA systems occurs during food-motivated learning, and demonstrate a capacity for bidirectional modulation of DA release in the mNAcSh by the MCH system.

## Introduction

The formation of associations between environmental cues and food availability is critical to survival. The brain integrates external sensory cues with information about internal states to drive appetition and consumption - ensuring the necessary intake of nutrients. In our current environment, however, associations between ubiquitous environmental cues (e.g., advertisements) and hedonically rewarding foods can drive overconsumption, leading to metabolic dysregulation and obesity, with attendant health consequences and costs [1].

The importance of the mesolimbic dopamine (DA) system in learning reward-predicting sensory cues is well established [2]. Increased DA release in the nucleus accumbens (NAc) signals the availability of rewards and stimulates food seeking [3,4]. The melanin-concentrating hormone (MCH) neuronal system, can increase food intake [5,6] and is known to interact with the mesolimbic DA system [7,8]. These neurons project widely throughout the brain, including to the ventral tegmental area (VTA) and NAc [9]. Interactions with reward systems have been postulated as a downstream mechanism underlying the effects of the MCH system on feeding and associative-learning [8,10,11]. MCH peptide injected into the NAc enhances food intake [12]. Optogenetic activation of MCH somas paired with food consumption enhances food intake [13], and/or increases preference for a paired food source while enhancing NAc DA tone [14]. Chemogenetic [15] or optogenetic activation of NAc-projecting MCH afferents [16] have similar behavioral effects. Studies have examined MCH-DA functional interactions, either in the context of consumption of drugs of abuse [7], in *ex vivo* preparations [17,18], or in feeding [19]. However, these studies generally focused exclusively on MCH peptide, as opposed to MCH neurons, and/or employed temporally or spatially imprecise methods.

Here, we used fiber-photometry to characterize MCH activity and DA release dynamics within the NAc with high temporal precision during feeding and Pavlovian conditioning. We observed that DA dynamics and MCH neuron activity are moderately correlated and the relationship between these systems shifts during “behaviorally engaged” states. Cue-aligned DA responses emerge over the course of Pavlovian conditioning, while MCH responses to both cue and food-reward appear early in training and remain consistent. Gain and loss-of function of the MCH system could bidirectionally modulate DA release. These results indicate that co-activation of these systems occurs during food-motivated learning, and demonstrate a capacity for bidirectional modulation of DA release in the NAc by the MCH system. These interactions likely serve to promote food consumption and shape the trajectory of associative-learning.

## Materials and Methods

### Mice

Protocols were approved by the University of Michigan’s Institutional Animal Care and Use Committee, and are in accordance with NIH guidelines for the use and care of Laboratory mice. MCH-Cre (Strain#:014099) mice and littermate controls were used in these experiments.

PmchΔVglut2 mice, Pmch-iCre crossed with Slc17a6tm1Lowl/J mice (JAX® stock # 012898) were also used (ref. PMID:17488640, PMID:39007235).

### *In vivo* fiber photometry

Photometry surgery and analysis were performed as in [20]; (though see *Supplementary methods*). For each recording session, the signal was converted to ΔF/F ((F – F0)/F0); where F0 was calculated as the 10th percentile of the entire fluorescence trace) and subsequently normalized as a z-scored ΔF/F.

### Feeding experiments

In ‘free feeding’ experiments, photometry and video recordings were initiated, and mice were placed in a fresh cage. Several pieces of regular chow were placed into one corner of the cage. Mice were then allowed to interact with the food for 20-30 minutes. For subsequent analyses, photometry data was synchronized with the video recording, and a human scorer noted the times during which the mouse consumed the chow.

### Pavlovian conditioning

Mice had two 30 min sessions to habituate to the recording chamber, in-cage FED3 [21], and optic cable and were allowed to freely receive pellets. Mice were trained across 3-11 subsequent sessions to associate a 1 kHz, 5s-duration tone with pellet delivery. ∼40 tones were played per session, with pellet delivery occurring 5 s after tone onset, and a variable 60-90s ITI.

### Gain and loss of function experiments

MCH-Cre mice were injected with AAV5-hSyn-DIO-HM4Di(Gi)-mcherry and dLight1.1 (mNAcSh). Prior to the start of 1-2 Pavlovian conditioning runs, mice were habituated to I.P. injections of saline. Mice were pretreated with either vehicle or 3mg/kg CNO 30 min. prior to running in the Pavlovian task.

MCH-Cre mice were injected with AAV5-Syn-FLEX-rc[ChrimsonR-tdTomato] and dLight1.1 (mNAcSh), and implanted with mNAcSh fibers. In a subset of trials (70%:30% stim:no-stim) either the tone, the pellet, or both were paired with optogenetic stimulation (5s-duration, 20Hz, 10ms pulse-width, 625 nm).

### Pharmacology

For MCHR1 inhibition experiments, 45 min. prior to the start of each run mice were treated with SNAP-94847 (25 mg/kg, I.P.).

### 2-color photometry analyses

Analysis was performed as in [22], using MATLAB scripts adapted from their work. Briefly, after normal signal processing, synchronously recorded GRAB-rDA1m and GCaMP were analyzed for cross-correlation, coherence and phase-offset. Coherograms used a multi-taper estimation with the chronux function cohgramc. To generate the phase-offset curves, we likewise adapted the analysis in [22] – briefly, the signals were processed with a Butterworth (bandpass) filter and the phase angle of the Hilbert function was extracted. Fluorescence amplitudes were extracted and averaged in 10 degree bins from -180:+180 degrees. The curves were normalized across the full oscillatory cycle and superimposed.

### Statistical analysis

Statistical analyses were performed using Prism 9.0 or Matlab software. Data presented met the assumptions of the statistical test employed. N numbers represent final numbers of healthy/validated mice, except in the case of one mouse where histology was unavailable.

## Results

### MCH and mesolimbic DA systems respond to discrete food-rewards, and are correlated during feeding

Cohorts of mice expressing the DA sensor dLight1.1, with recording fibers implanted in either the lateral core of the nucleus accumbens (lNAcc) or its medial shell region (mNAcSh) – where MCH+ fiber innervation is densest – were used to observe DA release dynamics. *pMCH-Cre* mice expressing GCaMP were used to observe somatic and axonal calcium-dynamics (Fig. 1A). We observed spontaneous dynamics in freely behaving mice (Fig. 1B). Both the MCH and DA systems were transiently activated by consumption of discrete food rewards (Fig. 1C) but were only minimally activated when a similarly-sized non-food object (a piece of bedding) was dropped into their cage.

**Figure 1.**
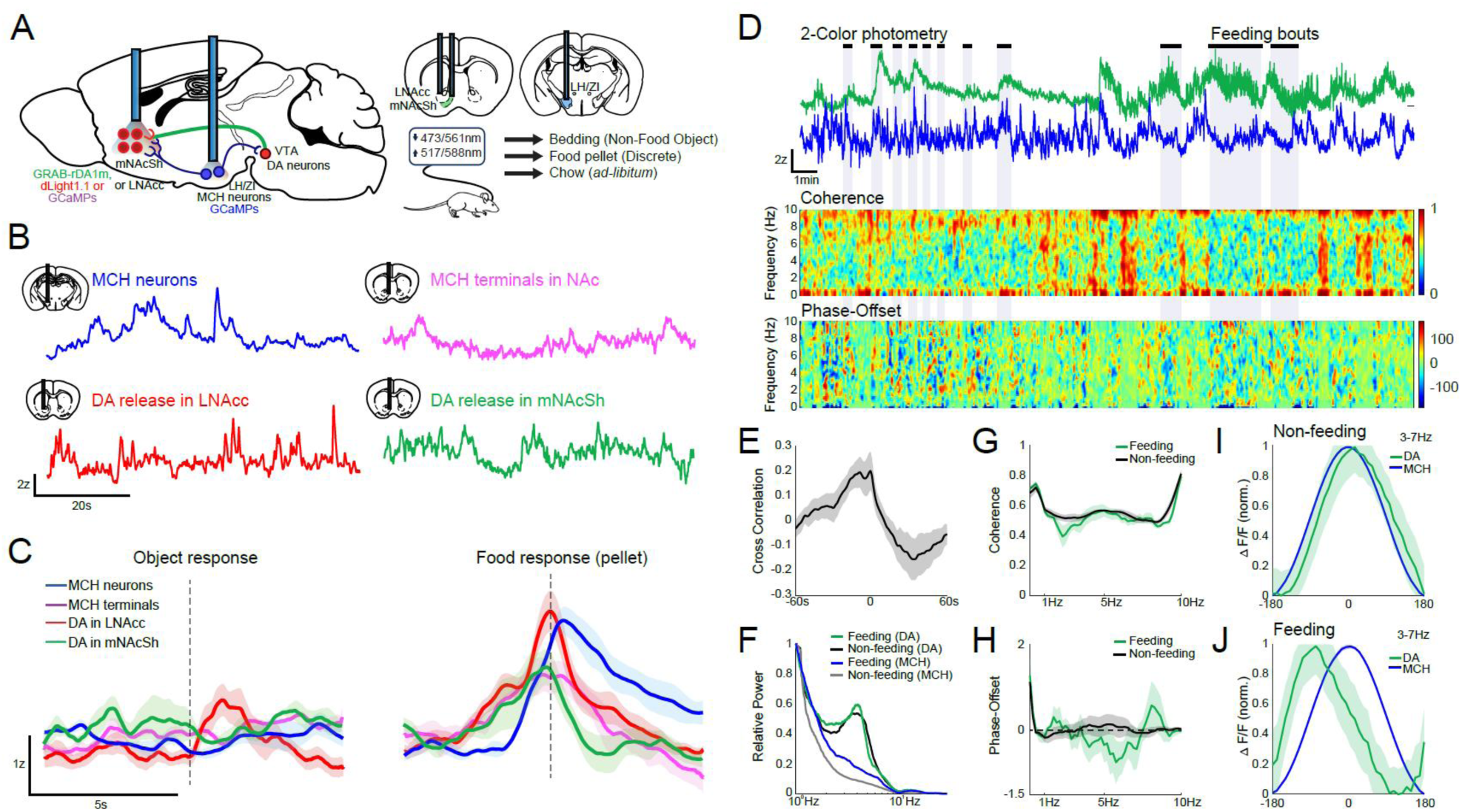
Fiber photometry recordings of DA release and MCH neuron activity during discrete and ad-libitum feeding. A) Schematic of experimental recordings. B) Example traces from MCH somas, MCH terminals in the mNAcSh, DA release dynamics in the lNAcc and mNAcSh. C) Example responses to pellet or control (bedding) availability. D) Simultaneous mNAcSh DA release dynamics (green) and MCH activity (blue) traces and feeding epochs during free-feeding. Coherogram and phase-offset relationship of 0.1-10Hz DA and MCH oscillations across the same recording. E) Mean cross- correlation between MCH and DA dynamics. F) Example Fast-Fourier Transform spectrum for 0.1-10Hz DA and MCH signals. G-H) Coherence and phase-offset from 0.1-10Hz during feeding and non-feeding epochs (n=4). I-J) Across-animals (n=4) phase-offset relationship in the 3-7Hz band during non-feeding and feeding epochs. See Fig. S1 for quantification.

Food-restricted mice were placed in a fresh cage while recording MCH/DA dynamics simultaneously, and videography was used to determine when mice were engaged in food consumption (Fig. 1C/D). Cross correlation analysis between MCH activity and DA release dynamics showed a moderate level of correlation over a range of several seconds (maximum correlation mean: 0.250±0.072) (Fig. 1E). We then refined our analysis to focus on a physiologically relevant range of oscillatory frequencies (i.e. 0.1-10Hz). As has been reported elsewhere [22], a prominent band of DA activity was observed around 4Hz (Fig. 1F). In comparison, MCH neurons did not show a clear peak of rhythmic activity (Fig. 1F). We found that MCH/DA coherence was moderate (∼0.6) in both feeding and non-feeding conditions (Fig. 1G). There was little offset between the two signals in non-feeding conditions (Fig. 1H/I); however, within the 3-7Hz band where DA showed rhythmic activity, DA release was observed to shift forward its phase relative with MCH activity during feeding (Fig. 1H/J, Fig. S1). This suggests that the two systems may be differentially driven during food consumption compared to non-consumption epochs.

### The MCH system responds to food-predictive cues early in Pavlovian conditioning

To have more control over the timing of feeding and to look at responses to food-predicting sensory cues, we shifted our focus to Pavlovian conditioning. Mice underwent between 3-11 days (7.2±0.63 runs/mouse) of conditioning with a daily run lasting 31-60 min (45.2±2.3 min/run). On average, mice took 37.5±2.5 (Range: 24-55) food pellets per run. We initially observed changes in lNAcc DA release dynamics over the course of Pavlovian conditioning (Fig. 2B). At early stages of conditioning, we observed a large phasic release peaking immediately prior to initial consumption of the food pellet, but no response to the auditory cue (Fig. 2C). After repeated pairings of the auditory cue with the pellet (44.0±4.3 trials/quintile, Range: 34-60), a strong phasic

**Figure 2.**
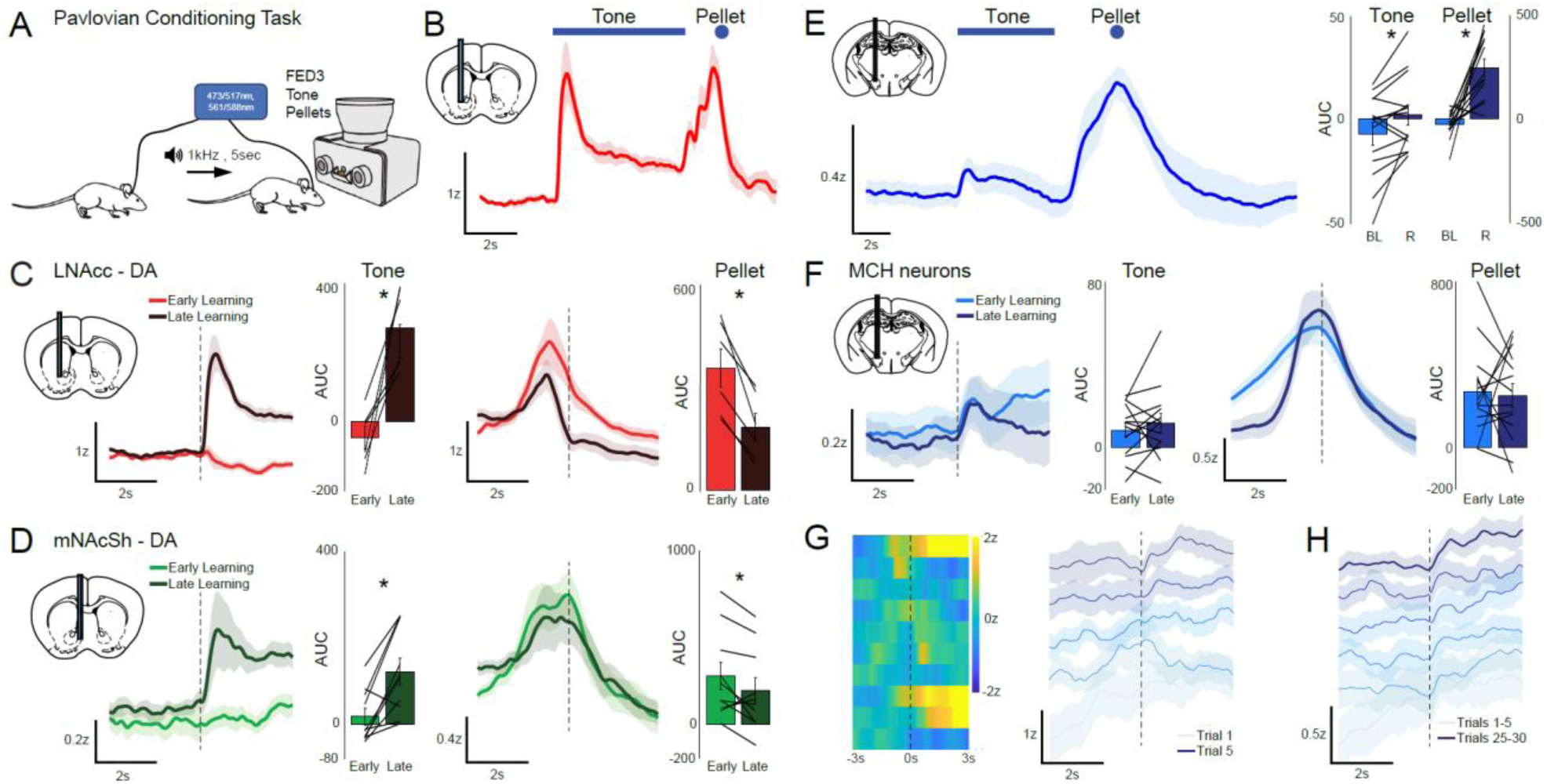
Changes in DA release dynamics and MCH neuron activity over Pavlovian conditioning. A) Schematic of Pavlovian conditioning. B) lNAcc DA release dynamics (n=6) showing response peaks to both the onset of the cue and the food-reward (pellet). C) lNAcc DA release (n=6) showing cue responses emerge over learning (inset: AUC of baseline-corrected Response Early vs. Late (Q1 vs. Q5), two-tailed paired t-test **p=0.0043, t=-4.945, df=5). Pellet responses diminish over learning (Early vs. Late, **p=0.0025, t=5.623, df=5). D) mNAcSh DA release (n=10) showing cue responses emerge over learning (Early vs. Late, two-tailed paired t-test **p=0.0024, t=-4.1737, df=9). Pellet responses diminish over learning (Early vs. Late, *p=0.0403, t=2.3933, df=9). E) MCH neuron activity (n=14) demonstrating responses to the cue and pellet (AUC of baseline vs. AUC of Response, all trials, tone *p=0.0273, t=-2.4856, df=13, pellet ***p=0.0001, t=-5.3403, df=13). F) MCH (n=14) cue and pellet responses do not change over learning once initially established (Early vs. Late, N.S.). G) Heatmap of peri-event (tone) MCH dynamics over the first 10 cue-reward pairings for a representative individual (left), and group mean traces (n=6), over the first 5 cue-reward pairings (right). H) Responses over the first 30 cue-reward pairings, after binning together every 5 consecutive trials (right). In responsive animals, MCH neuron tone- responses emerge rapidly, but are smaller when compared to MCH pellet responses or DA tone responses.

DA response to the cue developed, with the pellet response diminishing in magnitude. Over the course of a run in late learning, DA responses to the pellet but not the cue were attenuated – indicating some habituation to novelty or enhanced expectancy of the reward (Fig. S2B). These changes in lNAcc DA release dynamics are in line with classical reward-prediction error (RPE) [2]. In the mNAcSh, which is most strongly innervated by the MCH system, DA responses shifted from the pellet toward the cue in late-learning (Mean 46.9±5.9 trials/quintile, Range: 23.2-87) (Fig. 2D). DA release dynamics in the mNAcSh were qualitatively like those in the lNAcc but peaked during consumption rather than prior to consumption. DA release in the mNAcSh elicited by the cue in late learning tended to ‘ramp’ toward a plateau which was maintained through consumption of the pellet, as opposed to hitting a sharp peak followed by a rapid decline. Within a run, there was less decline in DA release in later vs. earlier trials (Fig. S2C).

MCH neurons were also activated strongly when mice approached and consumed the food pellets (Fig. 2E). Unlike DA release, there was a clearly defined but small (relative to the pellet response) response to the cue appearing very early in conditioning (Fig. 2E-H). This cue response was observable in a subset of animals (6/14) within the first 5 cue-reward pairings (Fig. 2G) and was consistent in shape and magnitude after that (Fig. 2H). There was little change in magnitude of the cue response from the first quintile of trials to the last (Fig. 2F) (59.4±4.1 trials/quintile, Range: 27.8-77.2), apart from a more consistent ‘ramping’ of activity toward the pellet response early on. There was no significant attenuation of the pellet response in late vs. early learning. We also confirmed that mNAcSh projecting MCH neurons are indeed activated during our Pavlovian conditioning task by recording from terminals in that region (Fig. S2A). Taken together, we can speculate that although the MCH system is activated by both the predictive cue and food reward across essentially all stages of conditioning, it does not function in an ‘RPE-like’ predictive manner – although there may also be a novelty component involved in MCH activation [23,24]. Co-activation of the MCH and DA systems across conditioning may shape behavioral responses, promoting task engagement and/or reward consumption and perhaps shaping activity in the NAc in both a pre- and post- synaptic manner by modulating DA release and interacting with postsynaptic medium-spiny neurons (MSNs) [11,19,25,26].

We performed within-animal simultaneous 2-color photometry of MCH activity and DA dynamics (Fig. S3A). We confirmed that, as suggested by our asynchronous observations, the MCH system is activated consistently by food-predictive cues very early in learning, prior to the emergence of DA responses to cue. This is clearest when observing responses during the first 30 trials (Fig. S3B-D). After roughly 50 trials in our paradigm, clear DA responses to cue emerged and continued to increase in magnitude, whereas MCH responses remained relatively small compared to pellet responses or DA responses (Fig. S3E).

### Correlation, coherence and phase relationships between MCH and DA systems during Pavlovian conditioning resemble feeding

We next examined the degree of coherence and the phase-relationship between activity in the MCH/DA systems over the course of Pavlovian learning. We compared activity within the intertrial- interval (ITI) to activity within a trial (i.e., during the cue-reward period; Fig. 3A). We observed a moderate degree of cross-correlation between MCH activity and DA release dynamics over a range of several seconds, with little change over learning (early learning maximum correlation mean = 0.292±0.066, late learning mean = 0.326±0.109). Similarly, we observed moderate coherence between the 2 signals in the 0.1-10Hz range during trials and in the ITI period (Fig. 3C/D). Unlike in the free-feeding paradigm, there was a clear phase-offset between the 2 signals in the 3-7Hz band whether we looked within the trial period or in the ITI (Fig. 3D/E, Fig. S1B/C). Activity during the Pavlovian conditioning paradigm therefore more closely resembled the feeding condition in this regard. This pattern of activity may therefore be reflective of task engagement, as opposed to a correlation with feeding *per se*.

**Figure 3.**
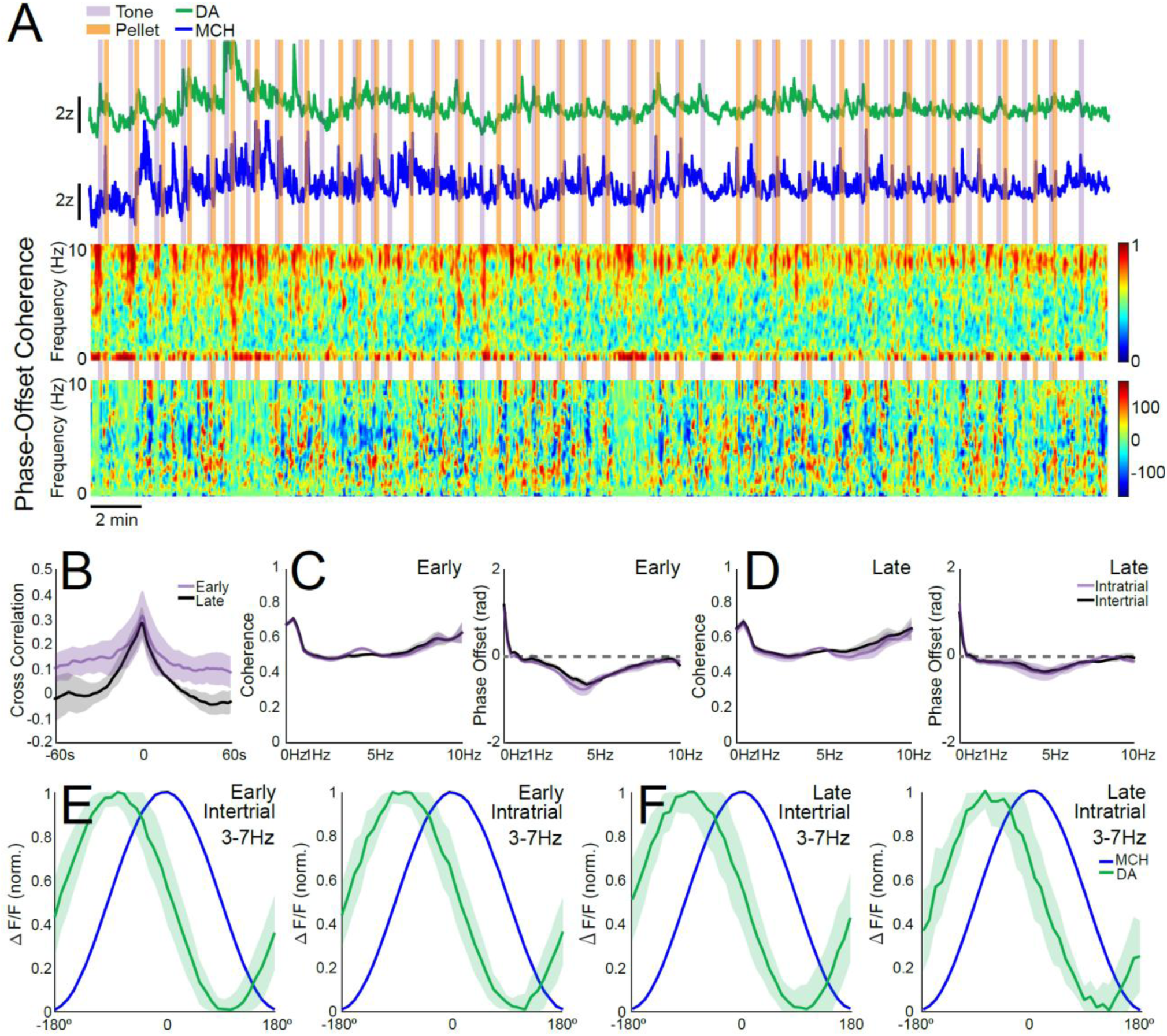
Coherence and phase-relationship between mNAcSh DA and LH/ZI MCH neuron dynamics across Pavlovian conditioning. A) Example simultaneous DA release and MCH activity during Pavlovian conditioning. Coherogram (middle, relative power) and phase-offset relationship (bottom, degrees) of 0.1-10Hz DA and MCH oscillations across the same recording. B) Cross-correlation between MCH/DA dynamics. C-D) Coherence between DA and MCH dynamics from 0.1-10Hz during cue/reward presentations or ITI, in early (n=3; C, Left) and late (n=3; D, Left) learning. Phase-offset relationship from 0.1-10Hz during cue/reward presentations or ITI, in early © and late (D) learning. E-F) Phase-offset relationship in the 3-7Hz band during ITI or cue/reward presentations, in early (E) and late (F) learning.

### Inhibition of the MCH system does not block the emergence of mNAcSh DA responses to cue in Pavlovian conditioning

To assess whether the MCH system is directly implicated in the emergence of DA responses to predictive cues in Pavlovian conditioning, we performed loss of function experiments on the MCH system. To constitutively block glutamate release from MCH neurons, we made use of *MCH- vglut2-floxed* mice. Blockade of glutamate release had little observable effect on the emergence of DA responses to auditory cues over learning (Fig. 4A). This suggests that glutamate release from MCH neurons is not critical for this type of learning. To block the effects of the MCH peptide, we pretreated experimental mice daily with the MCH-receptor 1 (MCHR1) antagonist SNAP- 94847. Pharmacological blockade of the MCHR1 did not completely prevent the emergence of DA responses to cue (Fig. 4B); however, it appeared to reduce task-engagement, which likely contributed to a delayed emergence of cue-response (Fig. S5).

**Figure 4.**
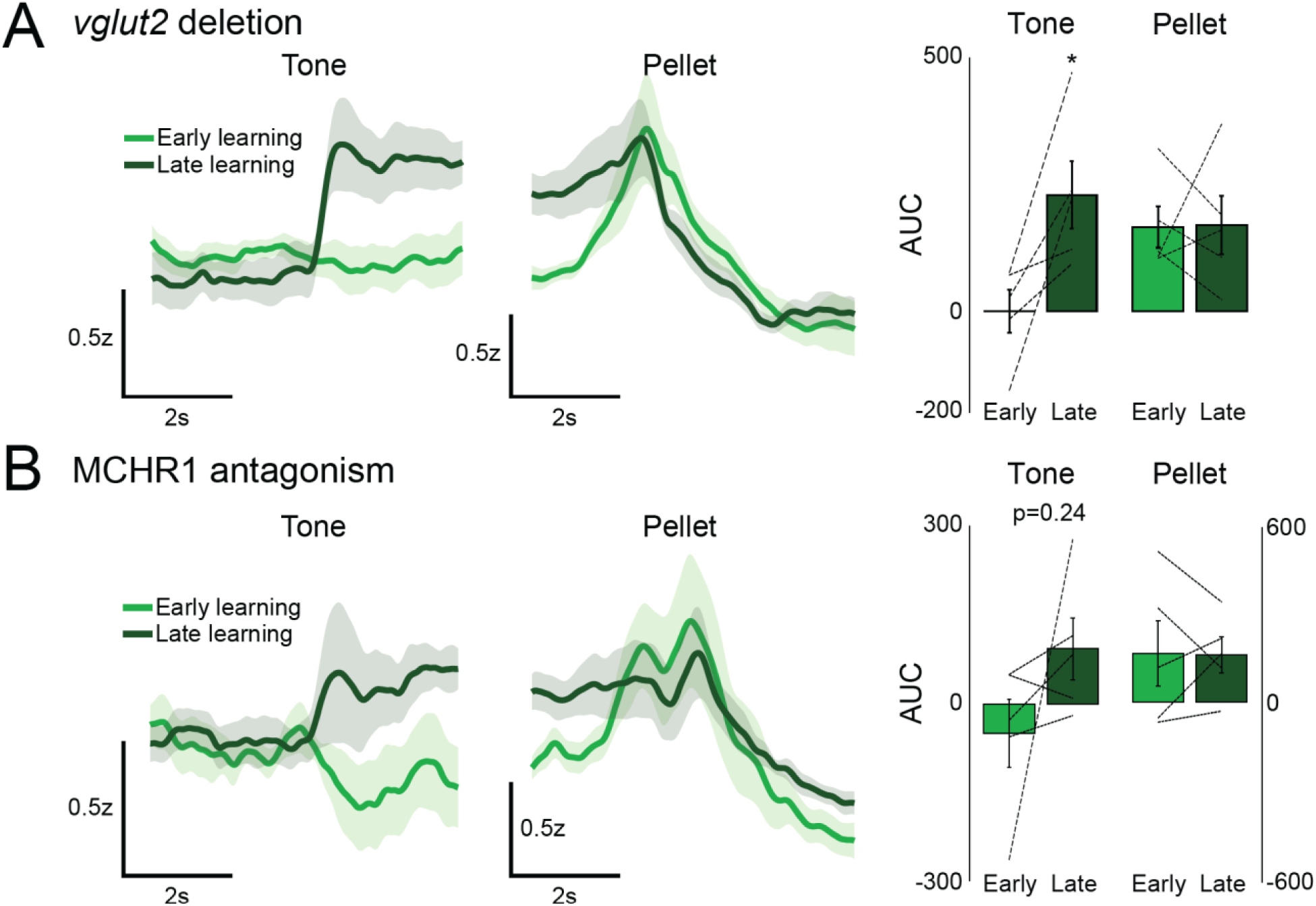
Effects of chronic MCH loss-of-function on mNAcSh DA release. Cue-aligned or pellet-aligned DA responses in MCH-vglut2-floxed mice during learning (AUC of Response – Baseline, Early vs. Late, n=5, tone two-tailed paired t-test *p=0.0298, t=-3.3052, df=4, pellet N.S.). B) Cue-aligned or pellet-aligned DA responses in MCHR1-inhibited mice during learning (Early vs. Late, n=5, tone two-tailed paired t-test p=0.2356, t=-1.3947, df=4, pellet NS).

### Acute manipulations of the MCH system demonstrate bidirectional modulation of DA release in the mNAcSh

Although the MCH system is not necessary for the emergence of normal DA dynamics in the mNAcSh over the course of Pavlovian conditioning, this does not preclude the possibility that the MCH system may modulate DA release in this context. In MCH-HM4Di mice, but not control mice, pretreatment with CNO led to a significant enhancement of DA release, particularly noticeable during the pellet response (Fig. 5A), compared to vehicle in the same animals. Likewise, when well-trained WT mice were pretreated with the MCHR1 antagonist, DA responses were significantly enhanced compared to vehicle in the same animals (Fig. 5B). These results are likely explained by ongoing (tonic) suppression of DA release by the MCH peptide. As these approaches are systemic it is possible that the effects are mediated at the level of the VTA or NAc.

**Figure 5.**
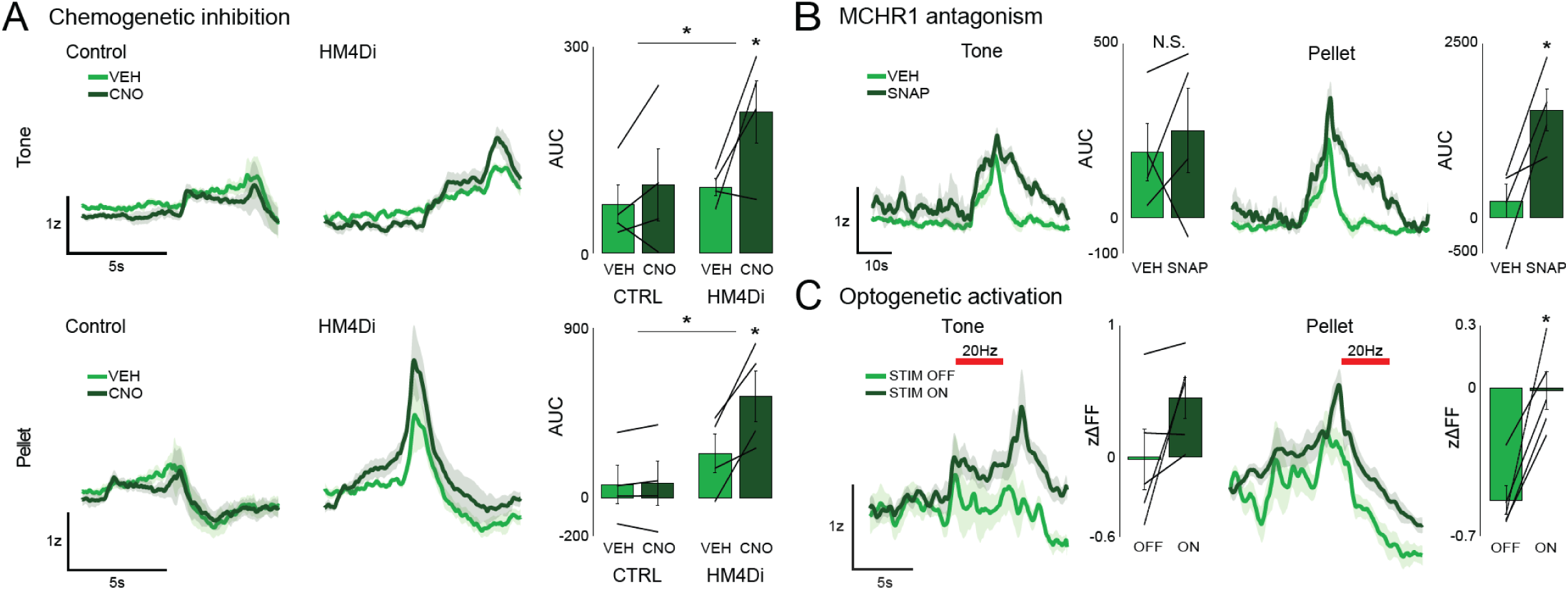
MCH neurons bidirectionally modulate DA release. A) Cue-aligned or pellet-aligned DA responses in VEH- or CNO- treated control (CTRL) or MCH- HM4Di animals during late-learning (n=4; 2-Way RM-ANOVA VEH vs CNO-treated, Control vs MCH-HM4Di group, Control n=4, HM4Di n=4. Tone: Significant overall effect of treatment *p=0.0135, treatment x genotype *p=0.0159, Sidak post-hoc, HM4Di-VEH vs. HM4Di-CNO *p=0.012. All other comparisons N.S. Pellet: VEH vs CNO-treated, Control vs MCH-HM4Di group, Control n=4, HM4Di n=4. Significant overall effect of treatment *p=0.018, treatment x genotype *p=0.049, Sidak post-hoc, HM4Di-VEH vs. HM4Di-CNO *p=0.0275. All other comparisons N.S.). Cue-aligned or pellet-aligned DA responses in animals treated with VEH or SNAP-94847 during late-learning (AUC of Response – Baseline, VEH vs. SNAP-treated, n=4, tone two-tailed paired t-test N.S., pellet *p=0.0317,t=-3.816, df=3). C) Cue-aligned or pellet-aligned DA responses in MCH-ChrimsonR mice, opto-paired (Opto) and unpaired trials (No Opto). Period of opto-excitation indicated in red (zΔFF over stim window, Opto unpaired (OFF) vs. paired (ON) trials, n=5, tone two-tailed paired t-test N.S., pellet **p=0.0051, t=-5.5734, df=4).

Enhancement of DA release appeared greater in the SNAP-94847 treated mice compared to chemo-inhibition of MCH neurons. We therefore hypothesized that glutamate release from MCH neurons might enhance DA transmission. Chemogenetic inhibition of MCH neurons is less effective at enhancing DA release versus MCHR1 blockade due to the opposing effects of inhibiting glutamate release from MCH neurons versus reducing MCH peptide release. To test this, we optogenetically stimulated MCH terminals in the mNAcSh. Once trained, we paired brief stimulation of MCH terminals with either the cue, the pellet, or both, with non-stimulated trials interspersed. We found that activation paired with either the tone or the pellet enhanced DA release compared to non-stimulated trials (Fig. 5C). Although it is possible MCH/MCHR1 signaling may play a role in this phasic enhancement, the relatively brief nature of the activation implies that it is more likely mediated by glutamate release.

## Discussion

Several studies on how the MCH system acts to influence hedonic feeding or food-motivated learning have pointed to an interaction between the MCH and the mesolimbic DA systems [11,12,14,16,17,19]. Here, we used recording and manipulation approaches to demonstrate the relationship between MCH activity and DA release with high levels of temporal and spatial precision, providing evidence of functional co-activation during feeding and learning, and of a causal interaction between MCH neuron activity and DA release in the mNAcSh.

We found that prior to any learning, MCH neurons are activated and DA release occurs in response to food-rewards. As in Subramanian et al. (2023) [10], we found that pairing an auditory cue with food delivery led to the emergence of MCH responses to the cue, indicating that they may signal reward expectancy. However, we did not see significant continued increases in magnitude of the MCH neuron response beyond the first day of learning. Rather, once cue-responses emerged, which occurred very early in training, they remained consistent over the course of training. This contrasts with the changes we observed in NAc DA release dynamics, which appeared later in training and continued to evolve and increase over the multi-day learning period. The early appearance of MCH cue-responses could suggest that MCH activity may shape emerging DA responses from very early in learning. MCH neuron activity has also been associated with novelty. Previous work [23,24] shows that MCH neurons are activated by novel-object presentation. Therefore, early cue-responses could signal novelty, except that we did not see attenuation of MCH cue responses across learning or within a run.

MCH and DA systems were co-activated during reward and cue presentation. Indeed, there is a relatively high degree of correlation and coherence between these two systems across multiple behavioral states (e.g., feeding, exploring, learning). This strengthens the argument that the systems are functionally coupled, and serve related behavioral functions, though these data do not identify the nature of this coupling. Additional studies may need to be conducted that employ better kinetically-matched sensors, longer recording periods, and more varied contexts (e.g., sated vs. food-deprived) to fully appreciate how these two systems are coupled.

We hypothesized that chronic loss of MCH neuronal function may slow the acquisition of DA responses to cues during learning. Neither blockade of glutamatergic signaling or MCH signaling abolished the normal DA dynamics over learning. Repeated MCHR1-antagonist treatment may, however, reduce task engagement, leading to slightly slower cue-response learning. These findings suggest that the MCH system is not required for the normal acquisition of phasic DA responses during Pavlovian conditioning, but instead may serve to augment food-seeking and drive task engagement. Both acute pharmacological inhibition of MCHR1 signaling and chemogenetic inhibition of MCH neurons produced increases in DA release. These data support previous work suggesting that MCH peptide has an inhibitory effect on DA signaling [25,26]. However, we also found that acute activation of MCH terminals in the mNAcSh increased DA release, suggesting that MCH neuronal activity can increase DA release despite possible release of MCH peptide. These observations may be reconciled by a proposed circuit in which MCH peptide/MCHR1 act tonically via Gi/o at VTA dopamine neurons to suppress DA tone in the NAc (Fig. S6) and glutamate signaling in the NAc (and potentially directly at VTA-DA) induces DA release. This proposed circuit fits with previous experiments [17,18,27,28] which found that VTA-

DA neurons were inhibited by MCH, in that *pMCH* deletion or MCR1 blockade led to a hyperdopaminergic state. Indeed, Spencer et al. (2024) [18] observed that most VTA-DA neurons express MCHR1 and superfusion of MCH suppressed firing in VTA-DA neurons in slice. Also of note, Scheeberger et al. (2018) [29] found that the effects of MCH neuron ablation on sucrose preference were primarily glutamate-mediated. Here, we add to our understanding of the MCH system and its role in modulating mesolimbic DA transmission in the context of Pavlovian learning and food consumption. MCH neurons are only one of many modulatory influences on reward systems that ultimately shape food-seeking and consummatory behavior.

## Acknowledgments

We thank members of the Burgess lab for helpful discussion. We thank the Elias lab for their contribution of pMCH-vglut2-floxed mice. This work was supported by a NIH 1F31NS132434 (BAT), Michigan Diabetes Research Center Pilot and Feasibility Award, a Whitehall Foundation new investigator grant, and 1R01DK129366 (CRB).

## Author contributions

LP, JRE, and JM contributed to the acquisition of data. LP, BAT, HCL, and CRB contributed to the analysis and interpretation of the data. LP and CRB contributed to drafting the manuscript.

## Supplementary methods

### Mice

Mice were housed in a University of Michigan vivarium in a temperature-controlled environment (12 h light and 12 h dark cycle; lights on at 2 AM). Following recovery from surgery, mice were food restricted as described below, but maintained *ad libitum* access to water. Protocols were approved by the University of Michigan’s Institutional Animal Care and Use Committee, and are in accordance with NIH guidelines for the use and care of Laboratory mice. MCH-Cre (Strain #: 014099) mice and littermate controls were used in these experiments. PmchΔVglut2 mice, Pmch-iCre crossed with Slc17a6tm1Lowl/J (Vglut2flox) mice (JAX® stock # 012898) were also used (ref. PMID: 17488640, PMID: 39007235).

### Surgery

Mice were deeply anesthetized by inhalation of 2% isoflurane and placed on a stereotaxic apparatus (Tujunga, CA). Following standard disinfection procedure, a small hole was drilled into the skull unilaterally at defined positions to target lNAcc (A/P: +1.2mm, M/L: - 1.3mm, D/V: -4.1mm), mNAcSh (A/P: +1.1mm, M/L: -0.53mm, D/V: -4.6mm), LH/ZI (A/P: - 1.6mm, M/L: +0.9mm, D/V: -4.8mm relative to bregma). A pulled-glass pipette was inserted into the brain, and virus was injected by picospritzer used to control injection speed at 25 nl/min. For fiber photometry experiments, 200 µL of AAV5-CAG-dLight1.1 (AddGene #111067-AAV5; titer 7 x 10^12^ genome copies per mL), AAV9-hSyn-GRAB-rDA1m (AddGene #140556-AAV9; titer 1 x 10^13^ genome copies per mL), or 500 µL bilaterally of AAV1.Syn.Flex.GCaMP6s.WPRE.SV40 (AddGene #100845-AAV1; titer 1 x 10^13^ genome copies per mL) or pGP-AAV-hSyn-FLEX-jGCaMP7s-WPRE (AddGene #104441-AAV9; titer 1 x 10^13^ genome copies per mL) was injected into the region of interest (Note: GCaMP mice were pooled regardless of virus used). For chemogenetic and optogenetic experiments, 500 µL of AAV5-hSyn-DIO-HM4Di(Gi)-mcherry (AddGene #44362-AAV5; titer 7 x 10^12^ genome copies per mL) or AAV5-Syn-FLEX-rc[ChrimsonR-tdTomato] (AddGene #6723- AAV5; titer 5 x 10^12^ genome copies per mL) were injected bilaterally in the LH region of *pMCH*-*Cre* mice, as well as AAV5-CAG-dLight1.1 in the striatum. An optic fiber (400-µm diameter core; BFH37-400 Multimode; NA 0.37 or 0.5; ThorLabs) was implanted over the region(s) of interest. Locations of optic fiber tips were identified based on the coordinates of Franklin and Paxinos [28]. After surgeries, mice were housed individually and monitored for proper recovery.

### *In vivo* fiber photometry

For behavioral experiments mice were food restricted to ∼85% body weight. Beginning 3 weeks after surgery, mice were connected to a fiber optic patch cable. Fiber optic patch cables (400 μm diameter; Doric Lenses) were firmly attached to the implanted fiber optic cannulae (Doric Lenses). LEDs (Plexon; 473 nm, Thorlabs; 405nm, Doric; 560nm) were set such that a light intensity of <0.2mW entered the brain; light intensity was kept constant across sessions for each mouse. Emission light was passed through a filter cube (Doric) before being focused onto a sensitive photodetector (Newport 2151 or Doric 119222-06). Signals were digitized at 1000 Hz using a National Instrument data acquisition system (or resampled up to 1000 Hz from 130 Hz in some experiments). In some experiments where 405 nm isosbestic signals were simultaneously acquired, the 405 nm excitation was either interleaved (time-domain) with the 473 nm excitation at 130Hz, or interleaved (frequency domain) at 217Hz or 319 Hz (10kHz recording). Multiplexed data was subsequently demodulated using custom MATLAB scripts. The signal was corrected by subtracting a double exponential fit, then adding back the mean of the trace. For each recording session, the signal was converted to ΔF/F ((F – F0)/F0); where F0 was calculated as the 10th percentile of the entire fluorescence trace) and subsequently normalized as a z-scored ΔF/F.

### Feeding experiments

Each session consisted of 4-10 alternating food and bedding drops. Only one session was run per day and mice underwent 1-2 sessions. All trials across days were pooled to calculate the response to food/bedding in each animal. In ‘free feeding’ experiments, photometry and video recordings were initiated, and mice were placed in a fresh cage. After habituation to the cage and establish baseline photometry levels, several pieces of regular chow were placed into one corner of the cage. Mice were then allowed to interact with the food for 20-30 minutes. For subsequent analyses, photometry data was synchronized with the video recording, and a human scorer noted the times during which the mouse consumed the chow.

### Pavlovian conditioning

Mice had two 30 min sessions to habituate to the recording chamber, in-cage FED3 [29], and optic cable and were allowed to freely receive pellets. Mice were trained across 3-11 subsequent sessions to associate a 1 kHz, 5s-duration tone with pellet delivery. ∼40 tones were played per session, with pellet delivery occurring 5 s after tone onset, and a variable 60-90s ITI.

### Gain and loss of function experiments

For DREADD experiments, MCH-Cre mice were injected with either AAV5-hSyn-DIO- HM4Di(Gi)-mcherry or AAV5-Syn-FLEX-rc[ChrimsonR-tdTomato] (for controls) in the LH/ZI and dLight1.1 (mNAcSh) during fiber implantation surgery. A minimum of 5 weeks was allowed for viral expression before experiments. Mice were run through the Pavlovian conditioning paradigm as described elsewhere in methods, except that (30 min.) prior to the start of 1-2 runs, mice were habituated to I.P. injections of saline. After mice were fully trained, they proceeded to the 2 DREADD-experimental days. On the first experimental day, mice were pretreated with vehicle (0.5% DMSO in 0.9% saline, I.P.) 30 min. prior to running in the Pavlovian task as before. On the 2^nd^ experimental day, mice were pretreated with 3mg/kg CNO (I.P.) dissolved in vehicle 30 min. prior to the task.

For optogenetics experiments, MCH-Cre mice were injected with AAV5-Syn-FLEX- rc[ChrimsonR-tdTomato] (in LH/ZI) and dLight1.1 (mNAcSh), and implanted with mNAcSh recording fibers. As elsewhere, mice had time to recover and undergo normal food- restriction and habituation. As in the DREADD experiments, mice were already trained on the Pavlovian conditioning paradigm prior to the onset of the optogenetic experiments. The optogenetic stimulation paradigm was identical to the normal Pavlovian conditioning paradigm, except that in a proportion of trials (70% :30% stim:no stim) either the tone, the pellet, or both were paired with optogenetic stimulation (5s-duration, 20Hz, 10ms pulse- width, 625 nm). Stimulation was delivered through the mNAcSh fiber, and was calibrated to approximately 9mW at the fiber tip using a portable light meter. Optogenetic stimulation was triggered by a custom Arduino script and was recorded with TTL pulses by the DAQ for subsequent analysis. Removal of stimulation artifacts proved unnecessary in these mice due to the frequency modulation of 473/405nm excitation light.

### Pharmacology

For the chronic MCHR1 inhibition experiment, dLight1.1-expressing mice were run through the Pavlovian conditioning paradigm as described; however, 45 min. prior to the start of each run, mice were treated with SNAP-94847 (25 mg/kg, I.P., dissolved in 20% w/v β- cyclodextrin/saline). This dose was chosen as a similar dose (30 mg/kg I.P.) has previously been reported to have behavioral effects and result in high MCHR1-occupancy (estimate 80%+) [30,31] in mice, and appeared to produce less sedation/behavioral inhibition compared to 30 mg/kg.

The acute MCHR1 inhibition experiment was conducted as above, except mice had been previously trained without pharmacology in the Pavlovian conditioning paradigm. As in the DREADD experiments, mice were initially pretreated with vehicle (20% w/v β- cyclodextrin/saline) on the first of 2 days. On the 2^nd^ experimental day, mice were pretreated with SNAP-94847 (25 mg/kg, I.P).

### Immunohistochemistry

Following the conclusion of all experiments, mice were anesthetized with pentobarbital and transcardially perfused with phosphate-buffered saline (PBS) followed by formalin. Once removed, brains were post-fixed in 10% formalin overnight then transferred to 20% PBS-sucrose for 48 hours. Brains were then frozen and 40μm coronal sections were cut using a freezing microtome.

For dLight immunohistochemistry, brain sections were washed in 0.1 M phosphate buffered saline pH 7.4, blocked in 3% normal donkey serum/0.25% Triton X100 in PBS for 1 h at room temperature and then incubated overnight at room temperature in blocking solution containing primary antiserum (rabbit anti-GFP, 488-conjugated, Novus #NB600- 308, 1:5000; rabbit anti-TH, Thermo Fisher Scientific #OPA1-04050, 1:5000). The next morning, sections were extensively washed in PBS before being mounted onto polarized slides.

Fluorescent images were captured with a Keyence BZ-X810 slide scanner microscope. All primary and secondary antibodies used are validated for species and application.

### 2-color photometry analyses

Analysis was performed as in [32], using MATLAB scripts adapted from their work. Briefly, after normal signal processing, synchronously recorded GRAB-rDA1m and GCaMP were analyzed for cross-correlation (using MATLAB – xcorr function), coherence and phase-offset in the 0-10Hz range. Coherograms used a multi-taper estimation with the chronux function cohgramc (window, 10 s; overlap, 5 s; step, 5; padding, 0). Feeding/Pavolvian epochs were expanded by 3s prior to their start and 5s at the end to catch transitions. To generate the phase-offset curves, we likewise adapted the analysis in [32] – briefly, the signals were processed with a Butterworth (bandpass) filter (Order 4, Range 1-3Hz or 3- 7Hz) and the phase angle of the Hilbert function (MATLAB - hilbert) was extracted.

Photometry fluorescence amplitudes were then extracted and averaged in 10 degree bins from -180 to +180 degrees. The resultant curves were normalized between 0 and 1 across the full oscillatory cycle and superimposed for the figures. Curves were smoothed with a moving average (2s - cross correlation, 0.6Hz - coherence/phase offset, or 60 degrees - offset curves) for figures to improve interpretability.

### Statistical analysis

Statistical analyses were performed using Prism 9.0 (GraphPad) or Matlab software. Data presented met the assumptions of the statistical test employed. Exclusion criteria for experimental mice were (i) sickness or death during the testing period (ii) if histological validation of the injection site demonstrated an absence of reporter gene expression (iii) or if histological validation demonstrated a mistargeted fiber placement. These criteria were established before data collection. N numbers represent final numbers of healthy/validated mice, except in the case of one mouse where histology was unavailable.

## Supplementary Figures and Legends

**Figure S1.**
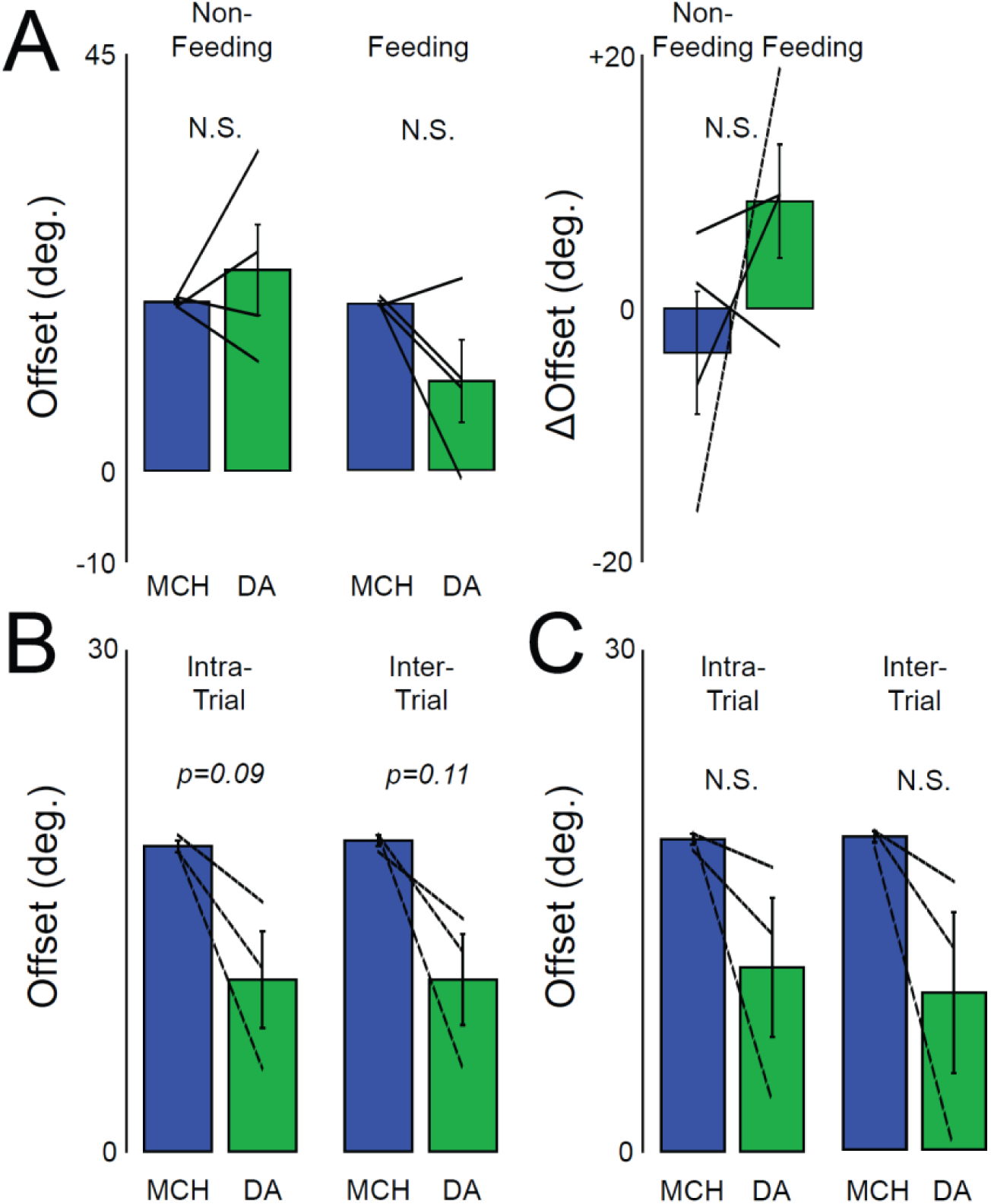
Quantification of phase-offset curves. **A)** Quantification of peak phase shift (degrees) of DA relative to MCH in non-feeding (two-tailed paired t-test p=0.5232, t=0.7207, df=3) (left) and feeding (two-tailed paired t-test p=0.15532, t=1.889, df=3) conditions, and the relative shifts (DA-MCH) in each state compared (right) (two-tailed paired t-test p=0.2616, t=1.380, df=3). **B)** Quantification of peak phase shift (degrees) of DA relative to MCH in intra-trial (two-tailed paired t-test p=0.0942, t=3.024, df=2) and inter-trial conditions (two-tailed paired t-test p=0.1066, t=2.813, df=2) (left) in early Pavlovian learning. **C)** Quantification of peak phase shift (degrees) of DA relative to MCH in intra-trial (two-tailed paired t-test p=0.2134, t=1.801, df=2) and inter-trial conditions (two-tailed paired t-test p=0.1729, t=2.081, df=2) (left) in late Pavlovian learning.

**Figure S2.**
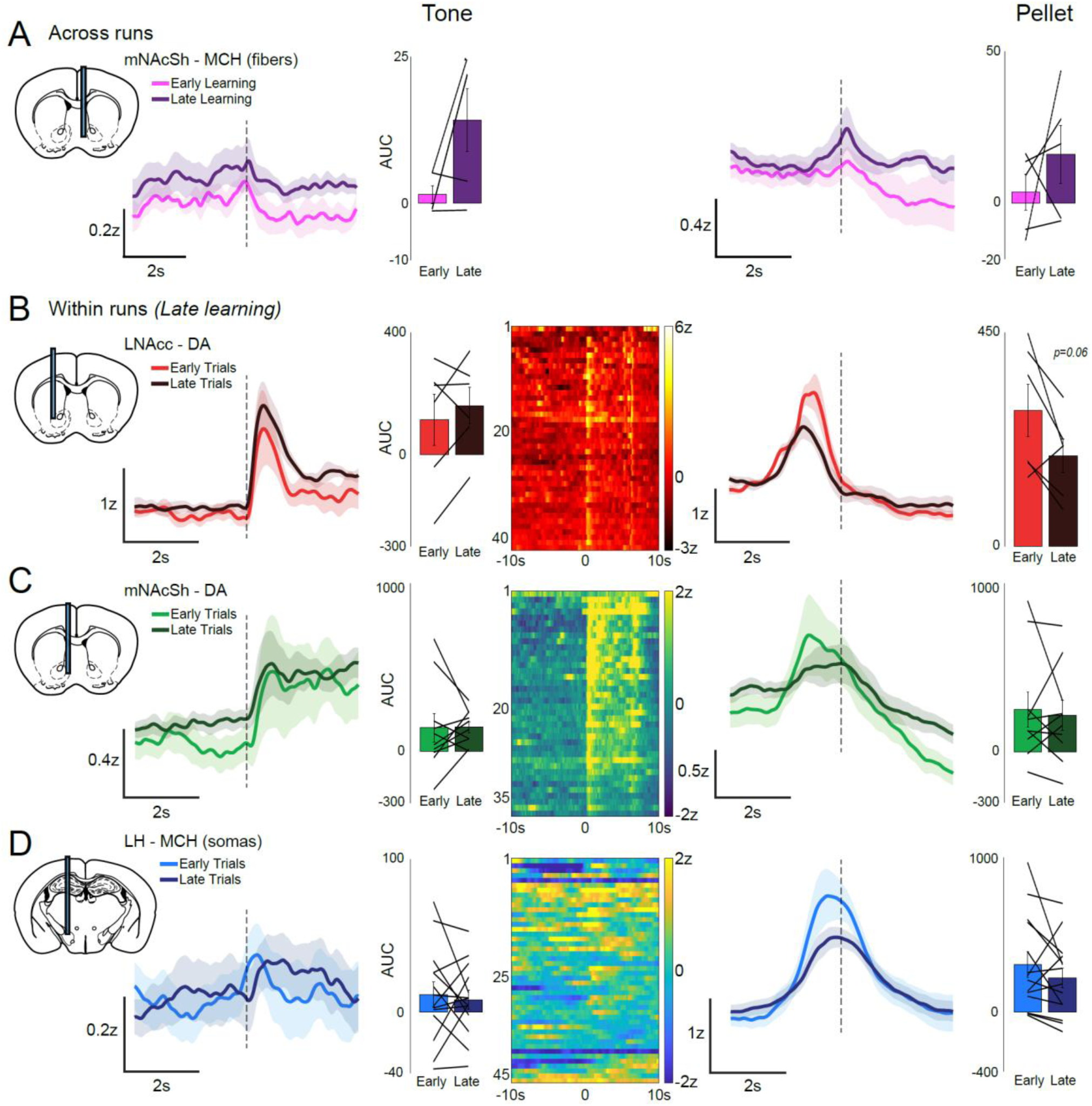
MCH axon-terminal responses to food-reward and cue in the mNAcSh, and changes in DA release dynamics and MCH neuron activity over the course of a Pavlovian conditioning run. A) mNAcSh MCH axon-terminal dynamics (mean across animals, n=5) during early and late learning. Cue responses trended towards an increase but generally showed low signal-to-noise ratio as well as an unstable baseline (tone) (AUC of Response – Baseline, Early vs. Late, n=5, two-tailed paired t-test p=0.0817, t=-2.3138, df=4). Food-reward (pellet) responses increase over learning N.S. (Early vs. Late, n=5, paired t-test=0.3819, t=-0.9815, df=4). Although recordings from terminals are often more technically challenging to achieve and are noisier (vs. somatic recordings), we were able to observe MCH axonal activation responses to both the food pellet and the cue in the mNAcSh, which were maintained across learning (Fig. S2A), and across a run. However, due to the lower signal quality of the mNAcSh terminal recordings, we focused on somatic recordings in subsequent experiments. B) lNAcc DA release dynamics in the early part of a run (first quintile of trials) vs the later part of a run (last quintile of trials) showing response to the cue (left) and acquisition of the food-reward (right). Inset: Quantifications of tone and pellet responses (AUC of baseline corrected Response, Early vs. Late trials, within-run, n=6, two-tailed paired t-test, tone: N.S.. pellet: p<0.0635, t=2.3753, df=5). Pellet responses, but not tone responses, are attenuated over the course of a run (adaptation). Heatmaps are from a representative individual, showing one run (43 trials), aligned to tone (left) or pellet (right). C) As in A, but for mNAcSh DA release dynamics. Inset: (AUC of Response – Baseline, Early vs. Late trials, within-run, n=9, two-tailed paired t-test, tone N.S. pellet N.S). Adaptation of pellet responses in the mNAcSh is less apparent vs. the lNAcc. Heatmap from a representative individual. D) As in A, but for MCH neuron dynamics. Inset: (AUC of Response – Baseline, Early vs. Late trials, within-run, n=9, two-tailed paired t-test, tone N.S. pellet N.S.). Heatmap from a representative individual.

**Figure S3.**
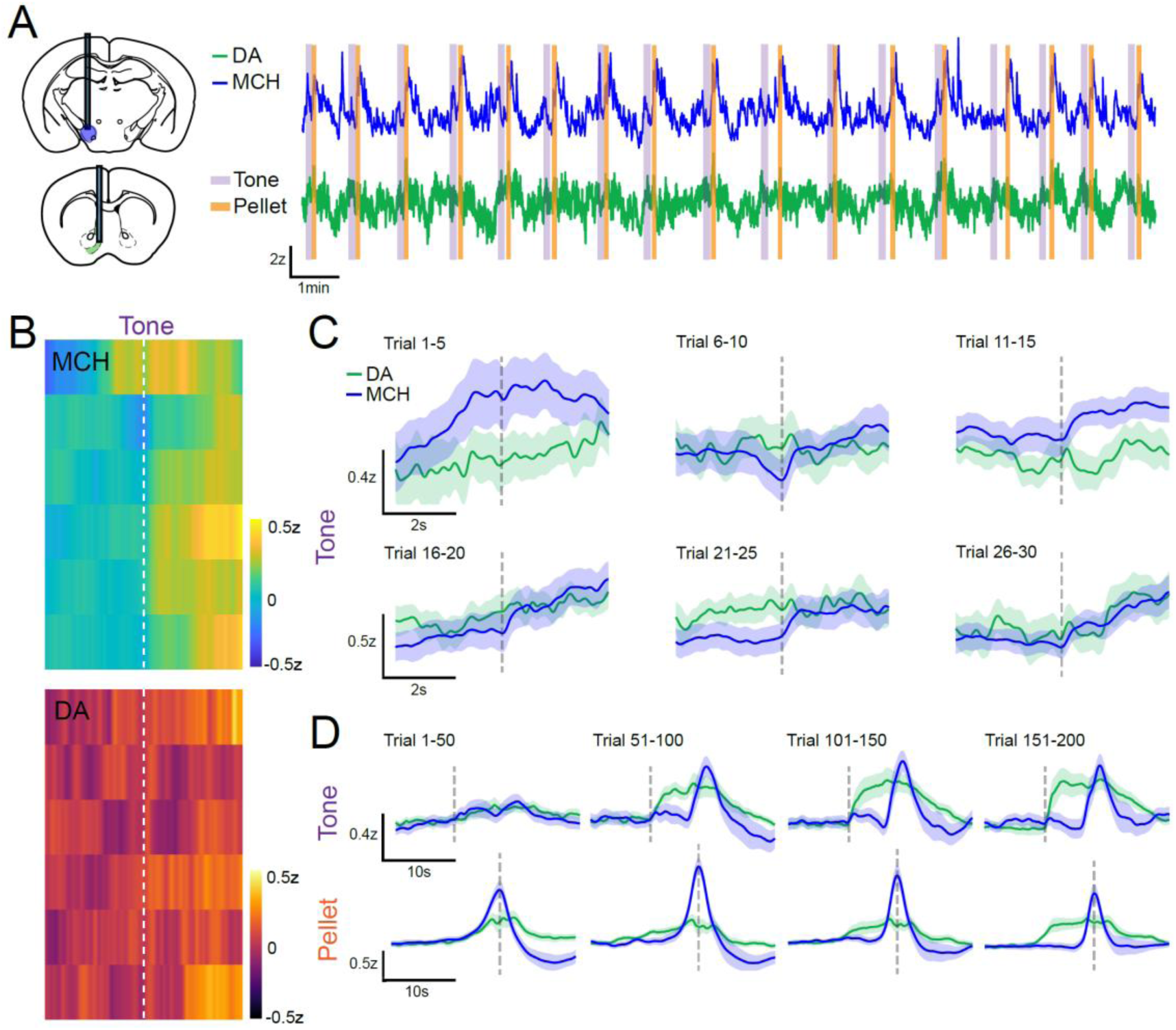
Simultaneous measurement of mNAcSh DA release and LH/ZI MCH neuron dynamics across Pavlovian conditioning. A) Example simultaneous traces of MCH neuron dynamics (blue, top) and mNAcSh DA release (red, bottom) during Pavlovian conditioning, with tone/pellet presentations demarcated. B) (Top) Example individual heatmap showing cue-aligned MCH responses over the first 30 cue-reward pairing, after binning together every 5 consecutive trials (n=6, ‘responders’ only). (Bottom) Example individual heatmap showing cue-aligned DA responses over the first 30 cue-reward pairing, after binning together every 5 consecutive trials (n=6, ‘responders’ only). C) Across-animals (n=6) mean cue (tone)-aligned MCH and DA responses over the first 30 cue-reward pairings, after binning together every 5 consecutive trials. A consistent but small MCH neuron response to tone emerges very early, prior to the DA tone response. D) Across-animals (n=6) mean cue (tone)-aligned (top) or pellet-aligned (bottom) MCH/DA responses over the first 200 cue-reward pairings, after binning together every 50 consecutive trials. Although the DA response to tone is of much greater magnitude, it emerges later than the MCH tone response.

**Figure S4.**
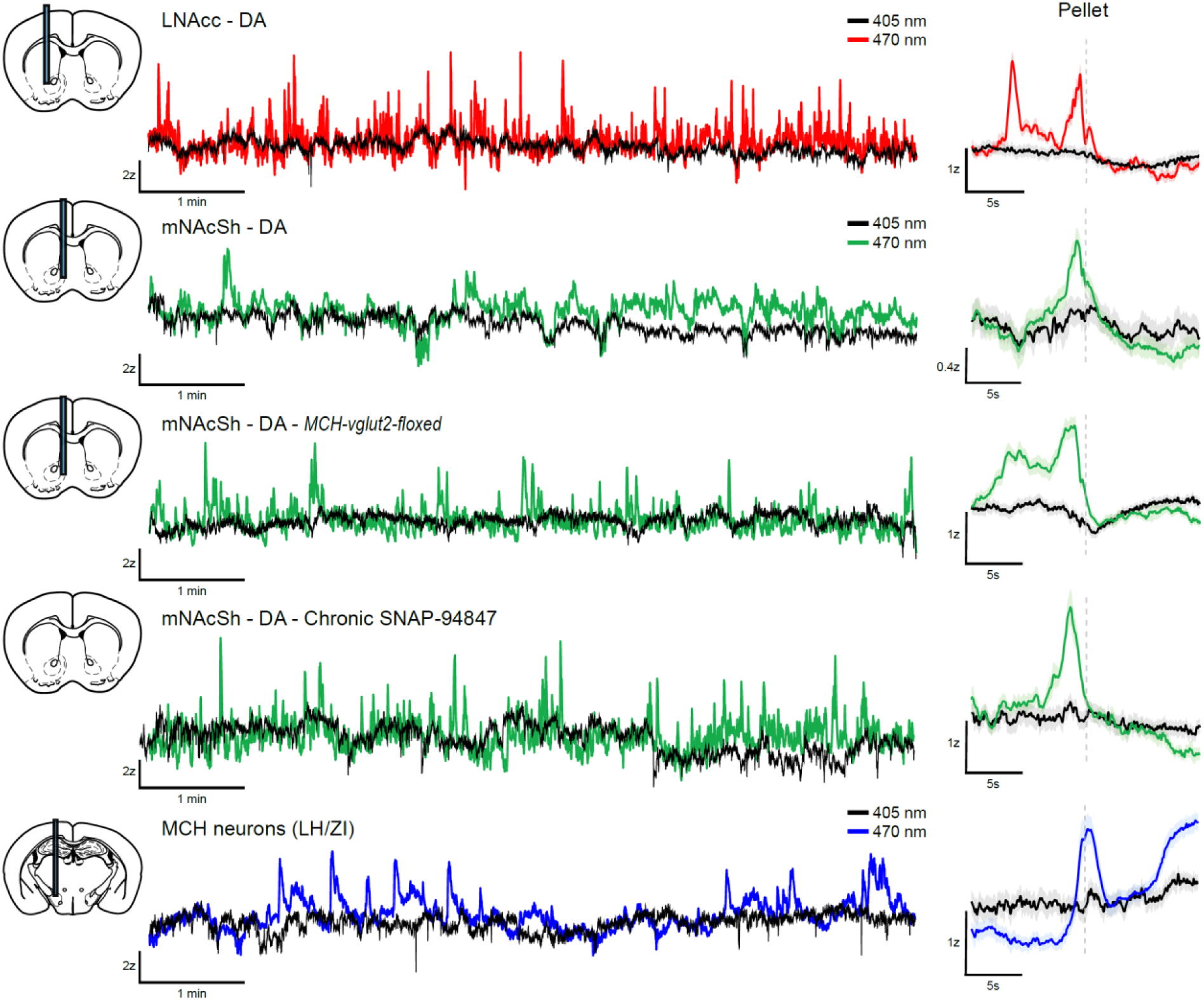
Example 405nm (isosbestic) traces vs. 470nm (calcium-dependent) traces during Pavlovian conditioning for various sensors and fiber locations. Individual traces of 470nm (calcium-dependent) and simultaneous 405nm (isosbestic) fluorescence during Pavlovian conditioning from the various sensors, fiber locations, and experiments (Left). Individual mean (single-run) peri-event 470nm and 405nm traces (time locked to pellet retrieval) from each sensor and location (Right). Some small motion effects are visible, but not prominent in averaged data – however, we lacked isosbestic data for many mice/groups.

**Figure S5.**
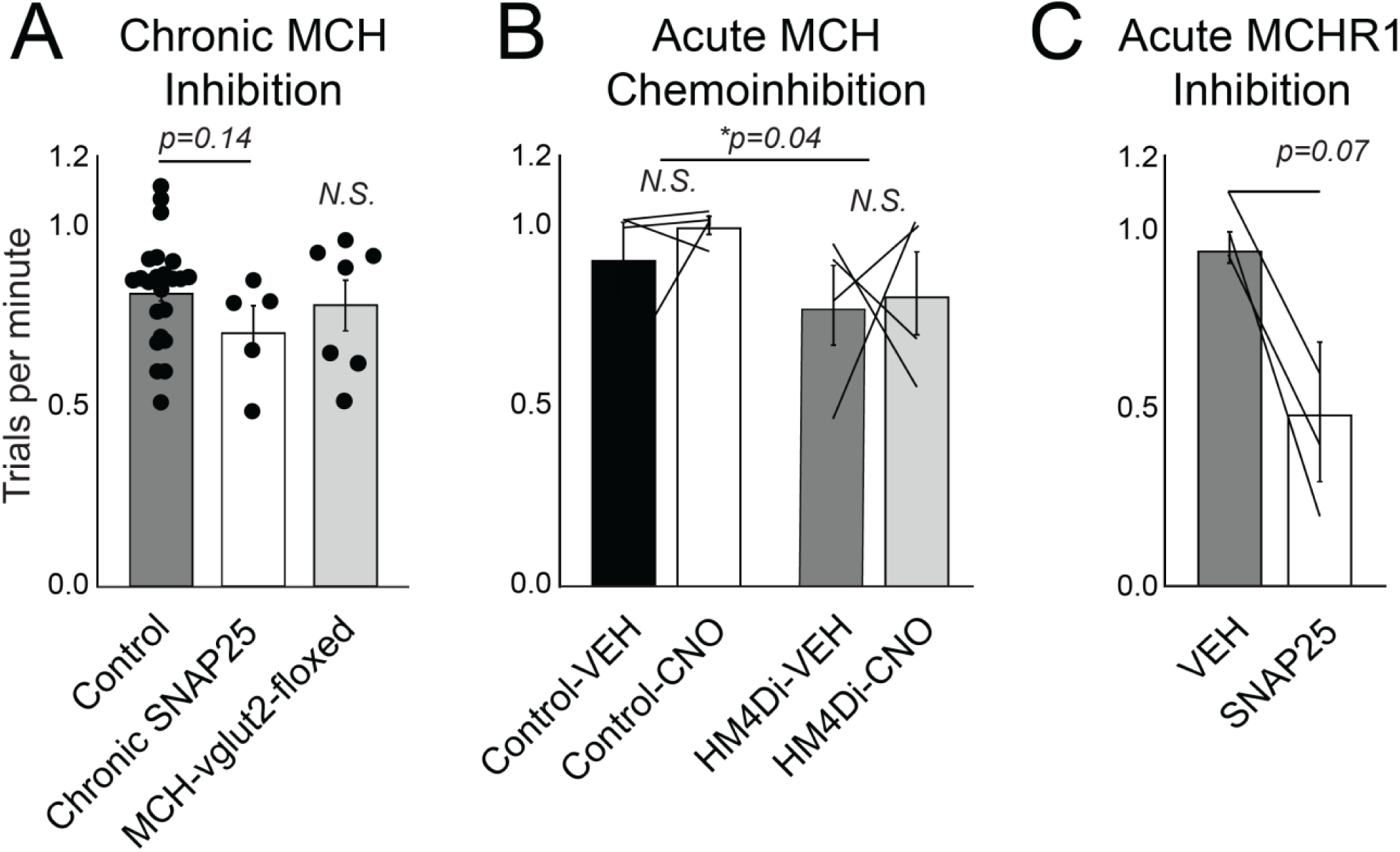
Effects of MCHR1/MCH neuronal inhibition on response rates in the Pavlovian conditioning task. A) Across-animals mean trial-rate (trials per minute) in chronic MCH system-inhibited animals. (Two tailed t-test vs. Control group, Control n=24, Chronic SNAP-94847 n=5, p=0.1371, t=1.532, df=27, MCH-vglut2-floxed n=5, N.S.) Although comparisons of fluorescence magnitude between disparate groups of animals with fiber photometry can be fraught, we also looked for fluorescence differences between treatment groups. Neither chronic loss-of-function manipulation produced statistically significant alterations in mean pellet response magnitude at any time point compared to controls. B) Across-animals mean trial-rate in acute MCH chemo-inhibited animals (late learning). (2- Way RmANOVA VEH vs CNO-treated, Control vs MCH-HM4Di group, Control n=4, HM4Di n=4, Significant overall effect of genotype p=0.0423, Treatment N.S., Treatment x Genotype N.S., all pairwise post-hocs, Sidak, N.S.) C) Mean trial-rate in acute MCHR1 inhibited animals (late learning). (Two-tailed paired t-test vehicle treated (VEH) vs SNAP-94847 (25mg/kg I.P.) treated (SNAP25), n=4, p=0.0702, t=2.759, df=3).

**Figure S6.**
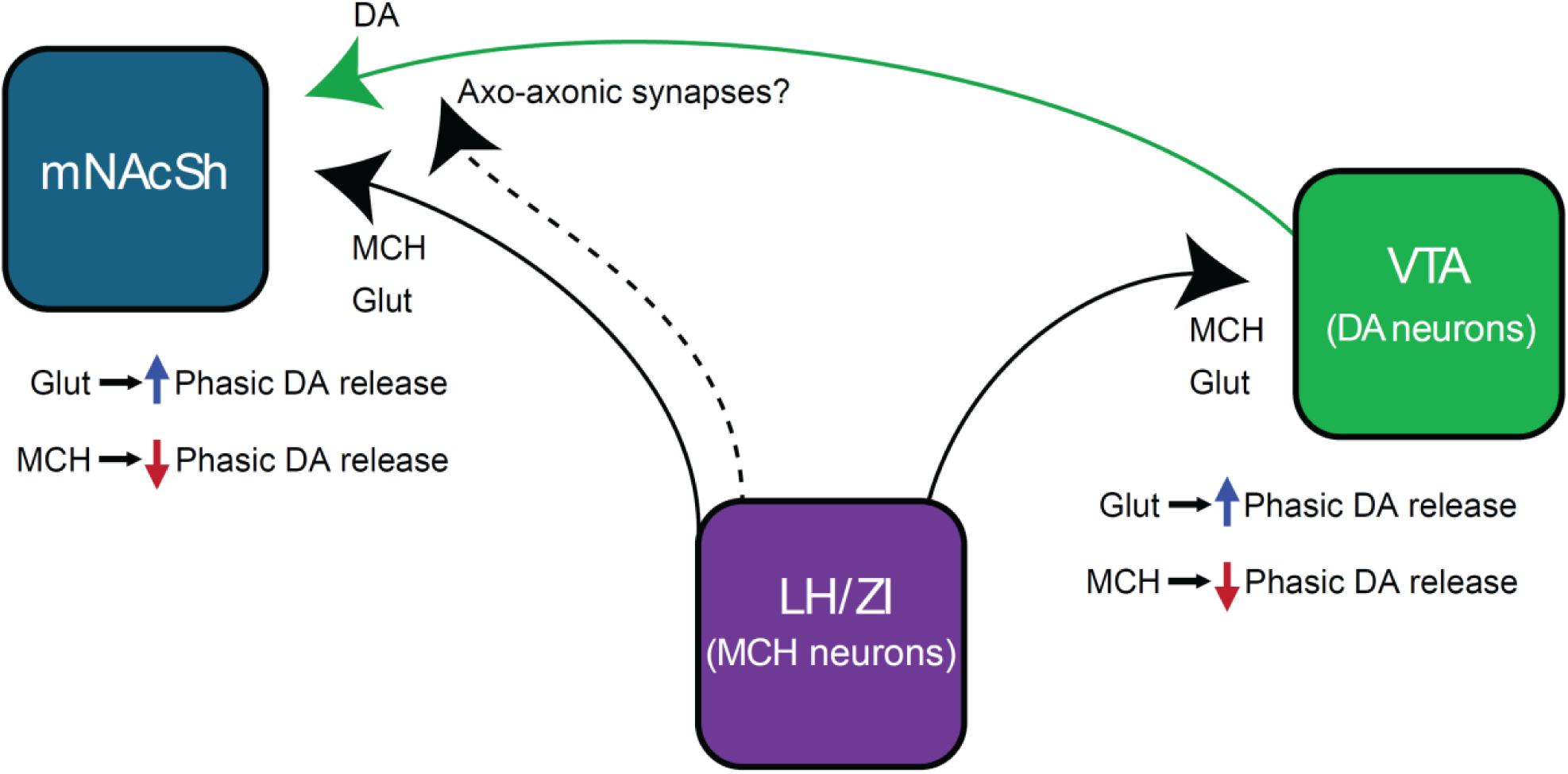
Effects of acute MCH gain-of-function on mNAcSh DA release dynamics in late-phase Pavlovian conditioning, and proposed circuit mechanisms.

**Figure S7.**
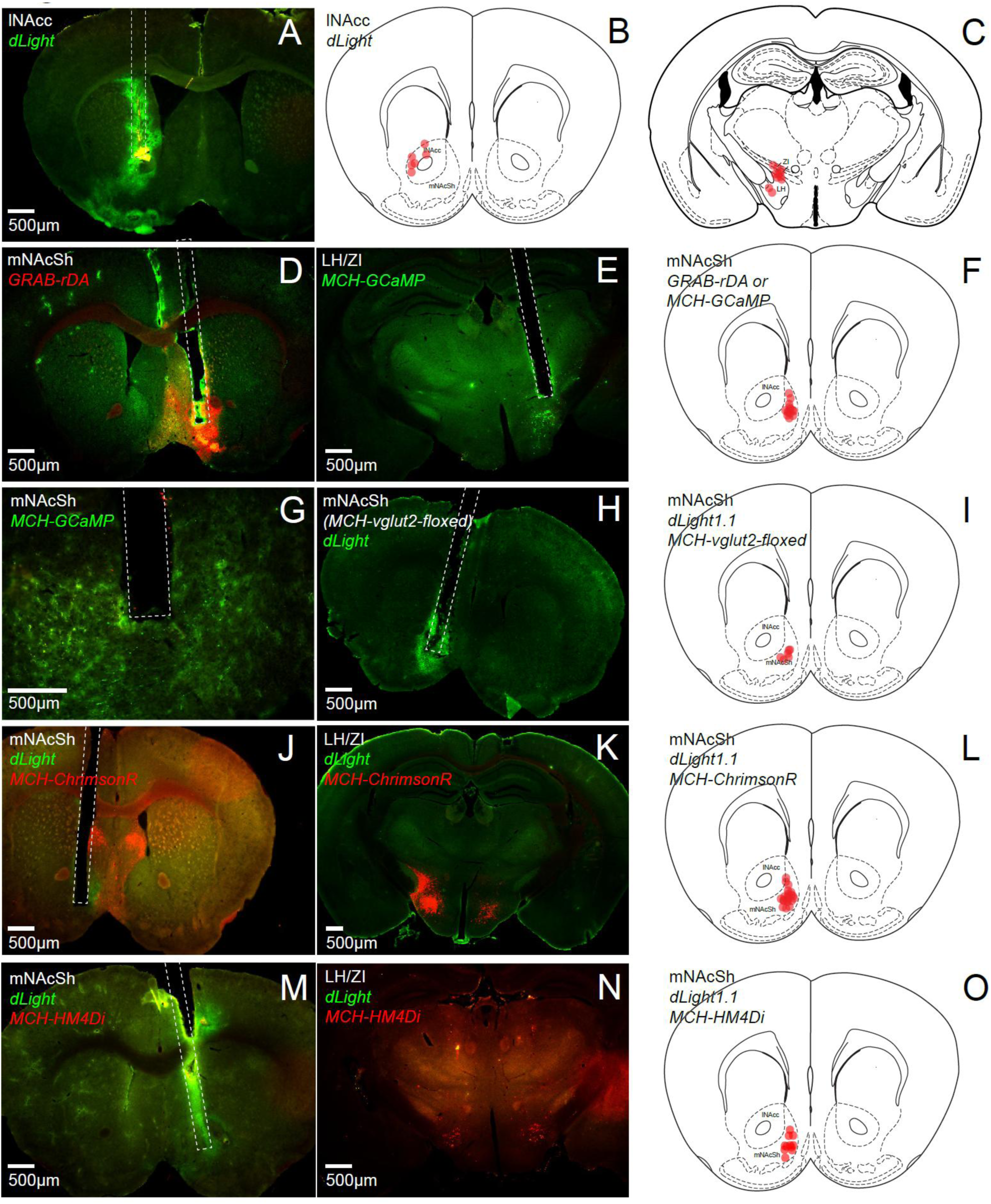
Sensor/actuator expression, fiber locations, and histology examples. A) Representative photomicrograph of dLight1.1 expression in lNAcc. (Fiber tract in hatched- white lines). B) Observed fiber-tip locations in experimental animals (lNAcc, dLight1.1). AP coordinate +1.1mm from bregma, may vary in individuals by approx. +/-0.2mm. C) Observed fiber-tip locations in experimental animals (LH/ZI, MCH-GCaMP). AP coordinate +1.7mm from bregma, may vary in individuals by approx. +/-0.2mm. D) Representative photomicrograph of GRAB-rDA1m expression in mNAcSh. E) Representative photomicrograph of MCH-GCaMP expression in LH/ZI. F) Observed fiber-tip locations in experimental animals (mNAcSh, GRAB-rDA1m). AP coordinate +1.1 mm from bregma, may vary in individuals by approx. +/-0.2mm. G) Close-up (20x objective) representative photomicrograph of MCH-GCaMP expression in mNAcSh. H) Representative photomicrograph of dLight1.1 expression in mNAcSh. I) Observed fiber-tip locations in experimental animals (mNAcSh, dLight1.1, MCH-vglut2- floxed). AP coordinate +1.1 mm from bregma, may vary in individuals by approx. +/-0.2mm. J) Representative photomicrograph of dLight1.1 expression in mNAcSh in MCH-ChrimsonR co-expressing mice. K) Representative photomicrograph of MCH-ChrimsonR expression in LH/ZI. L) Observed fiber-tip locations in experimental animals (mNAcSh, dLight1.1, MCH- ChrimsonR co-expressing mice). AP coordinate +1.1 mm from bregma, may vary in individuals by approx. +/-0.2mm. M) Representative photomicrograph of dLight1.1 expression in mNAcSh in MCH-HM4Di co-expressing mice. N) Representative photomicrograph of MCH-HM4Di expression in LH/ZI. O) Observed fiber-tip locations in experimental animals (mNAcSh, dLight1.1, MCH-HM4Di co-expressing mice). AP coordinate +1.1 mm from bregma, may vary in individuals by approx. +/- 0.2mm.

